# Treatment with both TGF-β1 and PDGF-BB disrupts the stiffness-dependent myofibroblast differentiation of corneal keratocytes

**DOI:** 10.1101/2024.02.29.582803

**Authors:** Krithika S. Iyer, Daniel P. Maruri, David W. Schmidtke, W. Matthew Petroll, Victor D. Varner

## Abstract

During corneal wound healing, stromal keratocytes transform into a repair phenotype that is driven by the release of cytokines, like transforming growth factor-beta 1 (TGF-β1) and platelet-derived growth factor-BB (PDGF-BB). Previous work has shown that TGF-β1 promotes the myofibroblast differentiation of corneal keratocytes in a manner that depends on PDGF signaling. In addition, changes in mechanical properties are known to regulate the TGF-β1-mediated differentiation of cultured keratocytes. While PDGF signaling acts synergistically with TGF-β1 during myofibroblast differentiation, how treatment with multiple growth factors affects stiffness-dependent differences in keratocyte behavior is unknown. Here, we treated primary corneal keratocytes with PDGF-BB and TGF-β1 and cultured them on polyacrylamide (PA) substrata of different stiffnesses. In the presence of TGF-β1 alone, the cells underwent stiffness-dependent myofibroblast differentiation. On stiff substrata, the cells developed robust stress fibers, exhibited high levels of ⍺-SMA staining, formed large focal adhesions (FAs), and exerted elevated contractile forces, whereas cells in a compliant microenvironment showed low levels of ⍺-SMA immunofluorescence, formed smaller focal adhesions, and exerted decreased contractile forces. When the cultured keratocytes were treated simultaneously with PDGF-BB however, increased levels of ⍺-SMA staining and stress fiber formation were observed on compliant substrata, even though the cells did not exhibit elevated contractility or focal adhesion size. Pharmacological inhibition of PDGF signaling disrupted the myofibroblast differentiation of cells cultured on substrata of all stiffnesses. These results indicate that treatment with PDGF-BB can decouple molecular markers of myofibroblast differentiation from the elevated contractile phenotype otherwise associated with these cells, suggesting that crosstalk in the mechanotransductive signaling pathways downstream of TGF-β1 and PDGF-BB can regulate the stiffness-dependent differentiation of cultured keratocytes.

**Statement of Significance:** In vitro experiments have shown that changes in ECM stiffness can regulate the differentiation of myofibroblasts. Typically, these assays involve the use of individual growth factors, but it is unclear how stiffness-dependent differences in cell behavior are affected by multiple cytokines. Here, we used primary corneal keratocytes to show that treatment with both TGF-β1 and PDGF-BB disrupts the dependency of myofibroblast differentiation on substratum stiffness. In the presence of both growth factors, keratocytes on soft substrates exhibited elevated ⍺-SMA immunofluorescence without a corresponding increase in contractility or focal adhesion formation. This result suggests that molecular markers of myofibroblast differentiation can be dissociated from the elevated contractile behavior associated with the myofibroblast phenotype, suggesting potential crosstalk in mechanotransductive signaling pathways downstream of TGF-β1 and PDGF-BB.

## Introduction

The cornea is the soft tissue at the anterior aspect of the eye, which is responsible for refracting light toward the retina (1). It consists of three cellular layers: the epithelium, stroma, and endothelium, but the stromal compartment accounts for the majority of the corneal thickness (2–4). The stroma contains a highly ordered ECM that is made up of lamellae of aligned collagen fibrils (2, 3, 5) with precise fibril diameter and spacing that allows light in the visible spectrum to pass freely through the tissue (6). A population of cells called corneal keratocytes populate the corneal stroma and are sandwiched between the collagen lamellae (1, 4). In their quiescent state, these cells maintain the microstructure of ECM and preserve corneal transparency (7–9), but following an injury, the keratocytes often are activated and transform into fibroblasts and myofibroblasts, which exert contractile forces and secrete fibrotic ECM that can disrupt the optical properties of the tissue and lead to corneal haze (10–12). Secreted growth factors, such as transforming growth factor-beta 1 (TGF-β1) and platelet derived growth factor-BB (PDGF-BB), among others, are released into the stromal space and have been shown to initiate this response (13–15). Among these soluble factors, TGF-β1 is thought to promote myofibroblast differentiation (16–18), while PDGF-BB stimulates keratocyte proliferation and motility (19). In addition, PDGF signaling has been shown to act synergistically with TGF-β1 to regulate the myofibroblast differentiation (20, 21).

Previous in vivo studies have also indicated that changes in mechanical properties accompany corneal wound healing (22, 23). Furthermore, substratum stiffness has been shown to influence the TGF-β1-induced myofibroblast differentiation of cultured corneal keratocytes (24, 25). In the presence of TGF-β1, cells in a compliant environment exhibit lower levels of myofibroblast differentiation, as indicated by fewer ⍺-SMA-positive cells (25, 26), as well as decreased cell-generated traction forces. In other experiments, changes in stiffness have been shown to impact PDGF-mediated keratocyte behavior, with cells exhibiting more elongated morphologies and proliferation at higher levels in stiffer microenvironments (27). Although PDGF signaling is known to influence TGF-β1-mediated myofibroblast differentiation (20, 21), it is unclear how changes in tissue mechanics impact the relative effects of TGF-β1 and PDGF on this process.

Here, we cultured corneal keratocytes in the presence of both TGF-β1 and PDGF-BB on functionalized 1 kPa, 10 kPa PA substrata and collagen-coated glass coverslips to study how changes in substratum stiffness affect ⍺-SMA expression, contractility, and the subcellular patterning of focal adhesions. These data were combined with pharmacological inhibition of PDGFR, to investigate if PDGF signaling contributes to stiffness-dependent differences in ⍺-SMA immunofluorescence.

## Methods

### Fabrication and functionalization of polyacrylamide (PA) gels

Polyacrylamide (PA) gels were fabricated on 30mm-diameter silanized glass coverslips, as described previously (25, 27, 28). Briefly, a 40 µL droplet of unpolymerized polyacrylamide solution was sandwiched between two surface-treated glass coverslips and allowed to polymerize for 30 minutes under vacuum. After the top slide had been carefully removed, the surface of the PA gel was functionalized using the heterobifunctional cross-linker sulfo-SANPAH (Pierce Biotechnology, Rockford, IL) and then treated with a 50 µg/ml solution of bovine type I collagen (PureCol; Advanced Biomatrix, San Diego, CA). PA gels of two different stiffnesses, 1 kPa and 10 kPa, approximating the stiffness of normal and fibrotic rabbit corneal tissue respectively (22, 23, 29) were fabricated. Collagen-coated glass coverslips were used as controls (25, 30, 31).

### Cell culture and reagents

As described previously, primary normal rabbit corneal keratocytes (NRKs) were harvested from New Zealand white rabbit eyes and plated on substrata of different stiffnesses (Pel Freez, Rogers, AR) (32–34). Briefly, the eyes were dissected in dishes containing RPMI-1640 medium supplemented with L-glutamine, sodium bicarbonate and 1% penicillin/streptomycin (Sigma-Aldrich, St. Louis, MO). The corneal epithelium was then removed by wiping the surface of the eye with an alcohol pad and scraping it with a surgical blade. Corneal buttons were then dissected using surgical scissors, and the endothelium was scraped away using a disposable scalpel. The dissected corneas were then incubated overnight at 37℃ in digestion medium containing 0.5 mg/ml hyaluronidase (Worthington Biochemicals, Lakewood, NJ), 2 mg/ml collagenase (Gibco-ThermoFisher, Waltham, MA), and 2% penicillin/streptomycin/amphotericin B (Lonza, Walkersville, MD). Afterward, the harvested NRKs were centrifuged-pelleted, resuspended, plated in T25 culture flasks, and cultured at 37℃ in serum-free conditions (i.e., DMEM supplemented with 1% RPMI vitamin mix (Sigma-Aldrich, St. Louis, MO), 100 μM nonessential amino acids (Invitrogen, Carlsbad, CA), 100 μg/ml ascorbic acid (Sigma-Aldrich, St. Louis, MO), and 1% penicillin/streptomycin/amphotericin B (26, 33)). First passage NRKs were used in all experiments, and the cells were plated on either collagen-functionalized PA gels or collagen-coated glass coverslips at 25,000 cells/ml and cultured in serum-free conditions for 24 hr to enable cell attachment. The culture medium was then replaced with medium containing either TGF-β1 (T7039; 5 ng/ml; Sigma-Aldrich, St. Louis, MO), PDGF-BB (PHG0045; 50 ng/ml; Gibco-ThermoFisher, Waltham, MA) or both (25, 27, 28, 35). The cells were then cultured at 37℃ for an additional 5 days and fixed. In some experiments, the PDGF receptor inhibitor, AG1296 (20μM; 502081527; Sigma-Aldrich, St. Louis, MO) was also added to the culture medium along with the exogenous growth factors (36). In all treatment conditions, a media swap was conducted after 48 hr of culture in medium.

### Fluorescence Microscopy

Cells were fixed in 3% paraformaldehyde in phosphate-buffered saline (PBS) for 15 minutes and washed 3 times with PBS. Fixed samples were then permeabilized in 0.5% Triton X-100 in PBS (PBS-T) for 30 min at room temperature and blocked with 1% bovine serum albumin fraction V (Equitech-Bio, Kerrville, TX) in PBS for 1 hr at room temperature. The samples were then washed three times in PBS and incubated with primary antibody for 2 hr at either room temperature, or at 37℃. The following primary antibodies were used: anti-⍺-smooth muscle actin (⍺-SMA) (A2547; 1:600; Sigma-Aldrich, St. Louis, MO), anti-phospho-myosin light chain (pMLC) (3671S; 1:200; Cell Signaling, Danvers, MA) and anti-vinculin (CP74; 1:600; MilliporeSigma, Burlington, MA). After washing, the cells were incubated with Alexa Fluor-conjugated secondary antibody (1:200; Invitrogen, Carlsbad, CA) and/or Alexa Fluor 594-conjugated phalloidin (1:200; Invitrogen, Carlsbad, CA) for 2 hr at room temperature on a shaker. The samples were then washed three times in PBS and incubated with 4’-6-diamidino-2-phenylindole (DAPI) (1:1000; Sigma-Aldrich) at room temperature for 20 min. Images of fixed samples were captured using a Zeiss LSM 800 laser scanning confocal microscope equipped with either a 20×, NA 0.80, EC Plan-Neofluar objective (Zeiss) or a 40x, NA 1.3, Plan-Apochromat objective (Zeiss) was used. Quantitative analysis was performed using ImageJ, as described previously (27, 28).

### Traction force microscopy

We used traction force microscopy (TFM) to quantify the mechanical stresses exerted by cultured NRKs (25, 28). As described previously, fluorescent polystyrene microspheres (FluoSpheres, 200 nm diameter, dark red; ThermoFisher) were suspended at a density of 0.04% in the unpolymerized PA solution before gel polymerization and PA substrata were fabricated as outlined above. These beads were used as fiducial markers to track the gel deformations generated by the cultured NRKs. The cells were plated in customized 6-well plates containing functionalized PA substrata of different stiffnesses and cultured in serum-free conditions for 24 hr to allow cell adhesion. The culture medium was then replaced with medium containing either TGF-β1, PDGF-BB or both growth factors, as described above, and cultured at 37℃ for an additional 2 days. We then performed a media swap and moved the cells to a humidified stage-top incubator on a Zeiss AxioObserver 7 microscope equipped with a motorized stage and an ApoTome.2 structured illumination module. Time-lapse phase contrast and epifluorescence images were captured for an additional 72 hours of culture at 1-hour intervals. At the end of the experiment, cells were lysed using a 5% solution of Triton X-100 in PBS, and a find set of images were acquired to capture the undeformed configuration of the gel. Cell-generated traction forces were then computed using the Particle Image Velocimetry (PIV) and Fourier Transform Traction Cytometry (FTTC) plugins in ImageJ (37, 38), as well as the measured mechanical properties of the gels (25).

### Statistical Analysis

Data represents mean ± standard deviation from at least three experimental replicates. Statistical comparisons were made in Prism 9 (GraphPad; San Diego, CA) using a two-way ANOVA followed by a Tukey post-hoc test with p-values specified in the figure captions. Relative frequency histograms were created in MATLAB and used to generate cumulative frequency plots, which were compared statistically using the Kolmogorov-Smirnov test within Prism 9.

## Results

### Keratocytes treated with both TGF-β1, and PDGF-BB exhibit elevated stress fiber formation and α-SMA staining even in a compliant microenvironment

Previous studies have shown that corneal keratocytes undergo stiffness-dependent differences in myofibroblast differentiation in response to TGF-β1 (24, 25, 28). It is thought that synergistic signaling between TGF-β1 and PDGF is important for myofibroblast differentiation (20, 21, 40), but it is unclear how treatment with PDGF affects stiffness-dependent TGF-β1-mediated keratocyte phenotypes. To answer this question, we cultured primary corneal keratocytes on soft (1 kPa) and stiff (10 kPa) PA substrata, as well as collagen-coated glass coverslips (∼GPa) in the presence of either TGF-β1, PDGF-BB, or both growth factors and quantified levels of ⍺-SMA staining and cytoskeletal organization (Fig. 1). The soft and stiff PA gels were selected since they approximate the stiffness of either normal or fibrotic rabbit corneal tissue (22, 23, 29); collagen-coated glass coverslips were used as controls, since they are the standard substratum for in vitro experiments involving corneal keratocytes (18, 21, 27, 30, 31). After 5 days of culture, NRKs maintained in serum-free medium retained a dendritic morphology on substrata of all stiffnesses and, consistent with previous work (25), did not form stress fibers nor stain positive for ⍺-SMA (Fig. 1A). In the presence of PDGF-BB, the keratocytes adopted a polarized morphology but did not exhibit detectable levels of ⍺-SMA immunofluorescence (Fig. 1B), though the cells did exhibit more elongated cell bodies on the stiffer substrata, as observed previously (27). When treated with TGF-β1 however, we observed stiffness-dependent myofibroblast differentiation, with clear differences in cell morphology, cytoskeletal organization, and ⍺-SMA staining (Fig. 1C). On stiff PA gels and collagen-coated glass coverslips, a larger fraction of NRKs formed stress fibers and exhibited ⍺-SMA immunofluorescence than observed among cells on soft PA gels. But when the keratocytes were cultured simultaneously with both PDGF-BB and TGF-β1, we no longer observed these stiffness-dependent differences in ⍺-SMA staining. Instead, NRKs cultured on soft PA gels had abundant stress fibers and exhibited the same elevated ⍺-SMA staining observed in stiffer microenvironments (Fig. 1D). In ease case, the cultured keratocytes exhibited a cell morphology that was hybrid of that observed in the presence of TGF-β1 or PDGF-BB alone: the central portion of the cell body was spread out, but each cell also extended thin elongated processes. In the presence of both growth factors, quantification across numerous experimental replicates indicated a significant increase in α-SMA staining and stress fiber formation among NRKs cultured on soft PA gels (Fig. 1E-F). In this compliant microenvironment, the cells also exhibited a larger cell area and length as compared to cells cultured with TGF-β1 alone (Fig. 1G-H), signifying the striking changes in morphology of corneal keratocytes due to the presence of both TGF-β1 and PDGF-BB.

**Figure 1.**
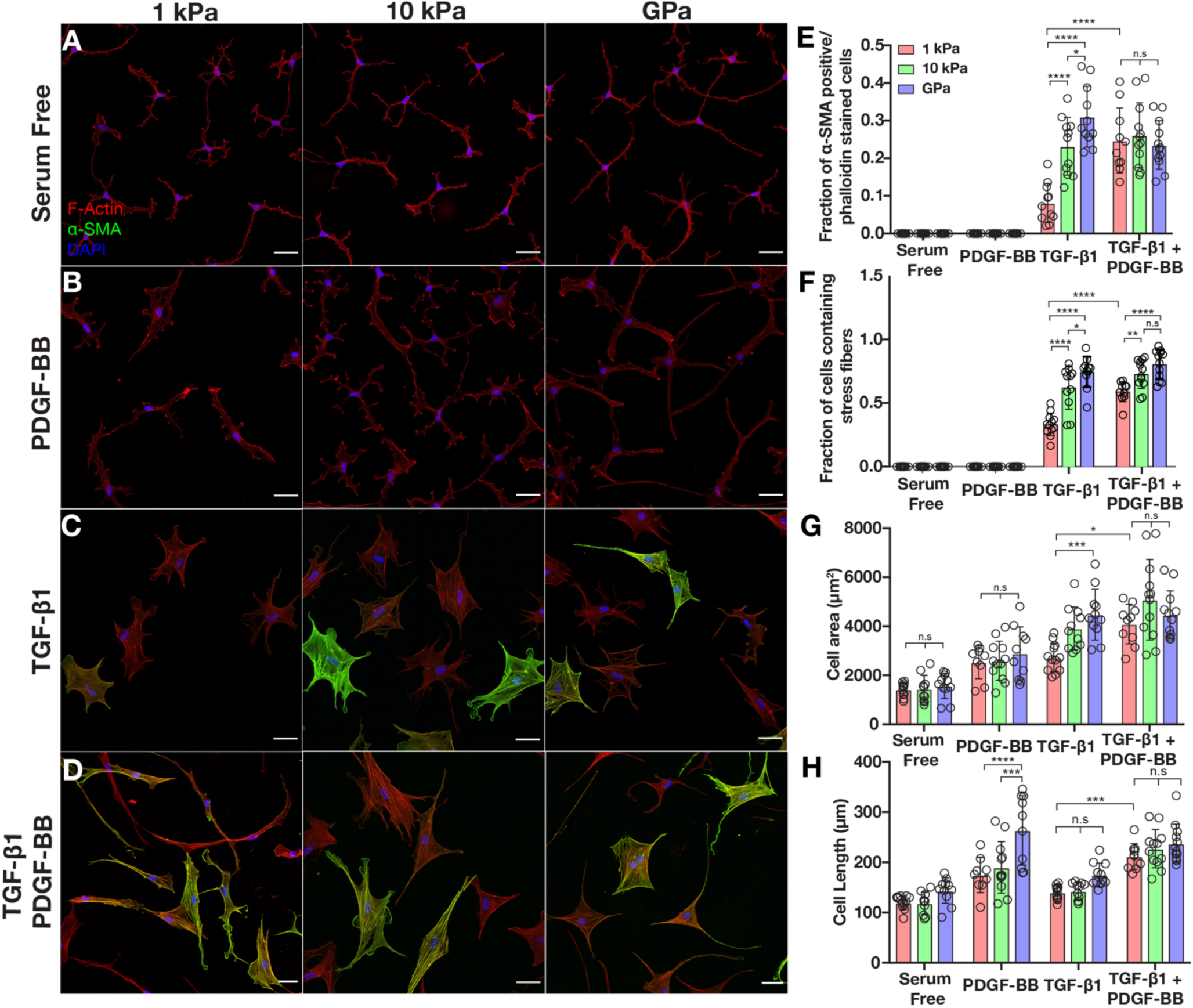
Expression of ⍺-SMA in the presence of TGF-β1 + PDGF-BB is independent of substratum stiffness. (A-D) Confocal images of phalloidin, α-SMA and DAPI immunofluorescence of corneal keratocytes in (A) serum free conditions or in the presence of (B) PDGF-BB (C) TGF-β1 or (D) TGF-β1 and PDGF-BB. Scale bar = 50 µm. (E-H) Quantification of the (E) fraction of α-SMA-positive cells, (F) fraction of cells containing stress fibers, (G) cell area and (H) cell length on substrata of varying stiffnesses in the presence of TGF-β1 and/or PDGF-BB. Error bars represent mean ± s.d. for n=12 substrates from 6 experimental replicates. A two-way ANOVA with a Tukey post-hoc test was used to evaluate significance among groups (*, p < 0.05; ****, p < 0.0001).

### Inhibition of PDGF signaling disrupts stiffness-dependent differences in myofibroblast differentiation

To confirm that signaling downstream of PDGF was driving these changes in keratocyte behavior, we repeated our PA gel experiments in the presence or absence of the PDGF receptor inhibitor AG1296. In serum-free conditions, the cells retained the same dendritic morphology even if PDGF signaling had been inhibited (Fig. 2A-B), and as expected, PDGF-BB-treated keratocytes no longer exhibited elongated cell morphologies when they were treated with AG1296 (Fig. 2C-D). Instead, the cells adopted dendritic geometries similar to those in serum-free conditions. In both cases, PDGFR inhibition did not alter ⍺-SMA staining, and no ⍺-SMA-positive cells were observed on substrata of any stiffness (Fig. 2A-D). In the presence of TGF-β1, however, PDGFR inhibition completely disrupted stiffness-dependent differences in ⍺-SMA immunofluorescence (Fig. 2E-F). In the presence of the inhibitor, most of the cells no longer formed stress fibers and exhibited dendritic cell morphologies. Similar changes in ⍺-SMA staining and cytoskeletal organization were observed among cells treated with both TGF-β1 and PDGF-BB, with fewer ⍺-SMA-positive cells on all substrata (Fig. 2G-H). Quantification across multiple experimental replicates revealed a significant decrease in stress fiber formation and ⍺-SMA immunofluorescence when cell cultured with either TGF-β1 alone, or both TGF-β1 and PDGF-BB (Fig. 2I-J), were treated with AG1296, suggesting that disrupted PDGF signaling suppresses myofibroblast differentiation on substrata of all stiffnesses.

**Figure 2.**
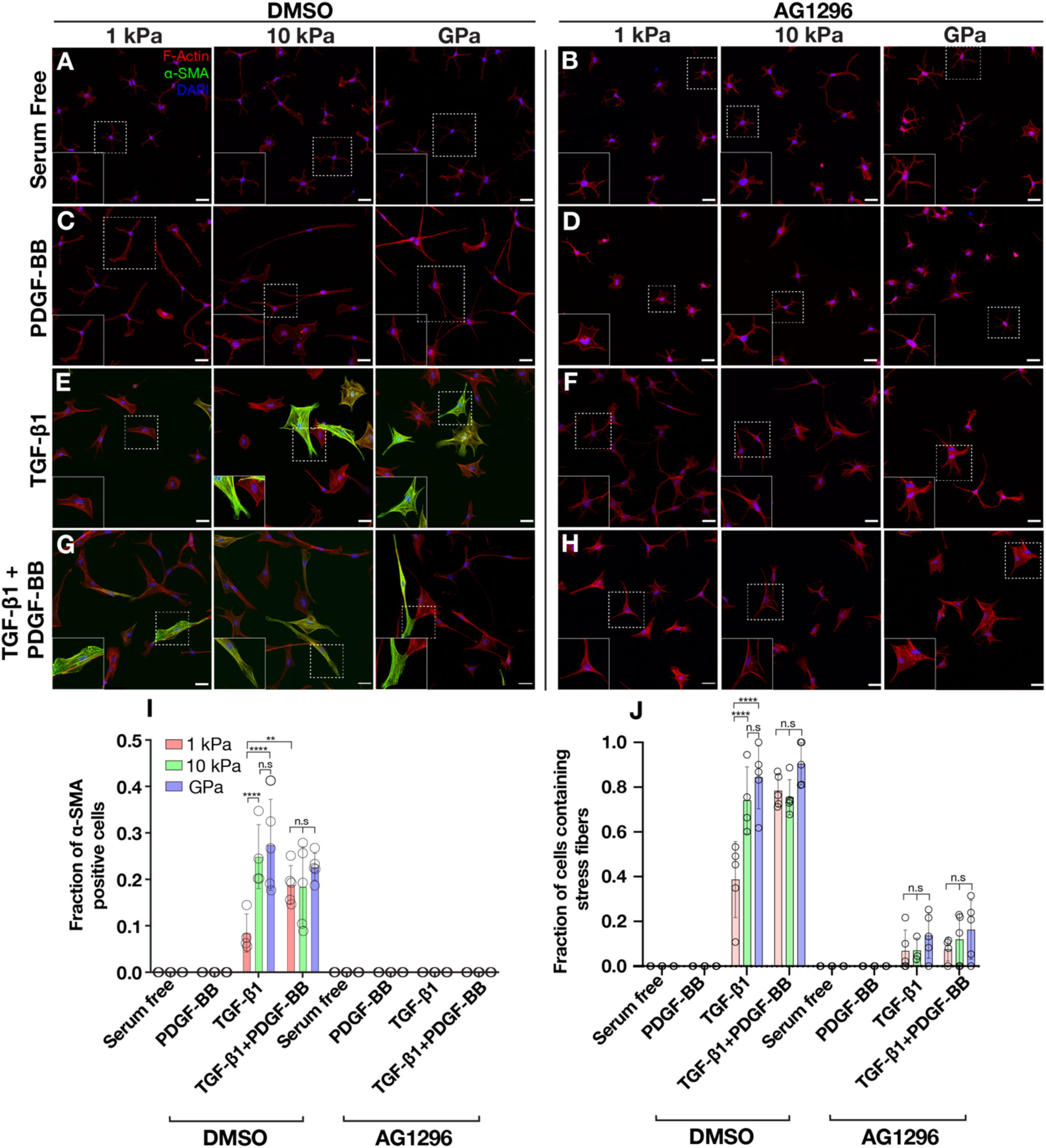
PDGF signaling is required for TGF-β1-induced myofibroblast differentiation. (A-H) Confocal images of phalloidin, α-SMA and DAPI immunofluorescence of corneal keratocytes in (A-B) serum free conditions or in the presence of (C-D) PDGF-BB (E-F) TGF-β1 or (G-H) TGF-β1 and PDGF-BB. Cells were cultured either in the presence of the (A, C, E, G) vehicle inhibitor, DMSO or (B, D, F, H) PDGF receptor inhibitor, AG1296. Scale bar = 50 µm. (I-J) Quantification of the (I) fraction of α-SMA-positive cells and (J) fraction of cells containing stress fibers on substrata of varying stiffnesses in the presence of TGF-β1 and/or PDGF-BB and/or AG1296. Error bars represent mean ± s.d. for n=5 substrates from 4 experimental replicates. A two-way ANOVA with a Tukey post-hoc test was used to evaluate significance among groups (*, p < 0.05; ****, p < 0.0001).

### Elevated contractility is decoupled from myofibroblast markers in the presence of TGF-β1 and PDGF-BB on soft substrata

Numerous studies have shown that myofibroblast differentiation is associated with increased cellular traction forces (41–43). To determine if elevated contractility is associated with increased DSMA staining among NRKs cultured on soft PA gels in the presence of both growth factors, we used TFM to quantify the traction stresses exerted by NRKs. Consistent with previous work in our lab (25, 28), TGF-β1-treated cells exhibited lower peak traction stresses when cultured on soft PA gels as opposed to stiff PA substrata (Fig. 3A). Contrastingly, in the presence of exogenous PDGF-BB, the NRKs exhibited extremely small traction stresses on substrata of all stiffnesses (Fig. S1). However, when treated with both TGF-β1 and PDGF-BB, the contractile behavior of the cells was similar to that observed with TGF-β1 alone (Fig. 3B). The cells exhibited stiffness-dependent differences in traction forces (Fig. 3C), reduced peak stresses observed on soft substrata, even though the NRKs exhibit elevated DSMA staining (Fig. 1D).

**Figure 3.**
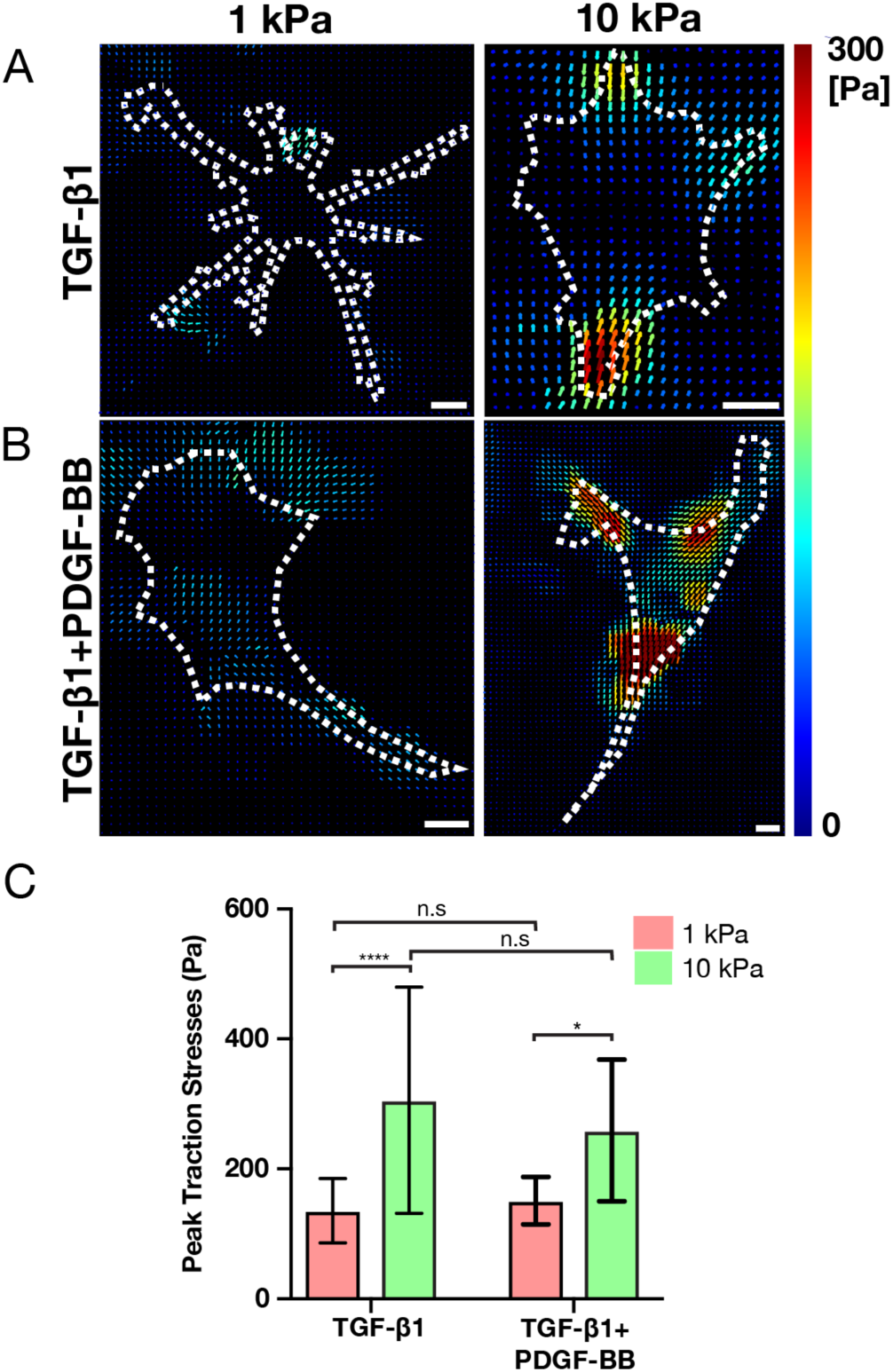
Traction stresses on compliant substrata do not increase in the presence of TGF-β1 and PDGF-BB compared to TGF-β1 alone. (A–B) Computed traction stresses for representative NRKs cultured on either 1 kPa or 10 kPa PA substrata, in the presence of (A) TGF-β1 only or (B) TGF-β1 and PDGF-BB. Scale bar = 20 µm. (C) Quantification of peak traction stress after 5 days of culture in the presence of growth factors. Error bars represent mean ± s.d. for individual cells from 4 experimental replicates. A two-way ANOVA with a Tukey post-hoc test was used to evaluate significance among groups (*, p < 0.05; ****, p < 0.0001).

We also stained the cells for pMLC immunofluorescence to visualize subcellular regions of active actomyosin contractility. As described previously (25, 28), keratocytes treated with TGF-β1 showed clear stiffness-dependent differences in pMLC localization (Fig. 4A-C). On soft PA gels, pMLC immunofluorescence was localized primarily at the tips of thin cellular extensions (Fig. 4A), while on stiffer PA gels and collagen-coated glass coverslips, pMLC colocalized with the numerous stress fibers that spanned the cell body (Fig. 4B-C). In the presence of both TGF-D1 and PDGF-BB, we observed similar stiffness-dependent differences in pMLC localization (Fig. 4D-F). As observed in TGF-β1 alone, pMLC colocalized with stress fibers in cells in stiff microenvironments (Fig. 4E-F), but on soft PA substrata, even when cells exhibit both stress fiber and ⍺-SMA staining, the pMLC immunofluorescence was sequestered primarily to the tips of thin cell extensions and did not colocalize with stress fibers (Fig. 4D). These data are in accord with our traction force measurements and suggest the presence TGF-β1 and PDGF-BB, in a compliant microenvironment, decouple ⍺-SMA staining from the elevated contractile phenotype typically associated with myofibroblast differentiation.

**Figure 4.**
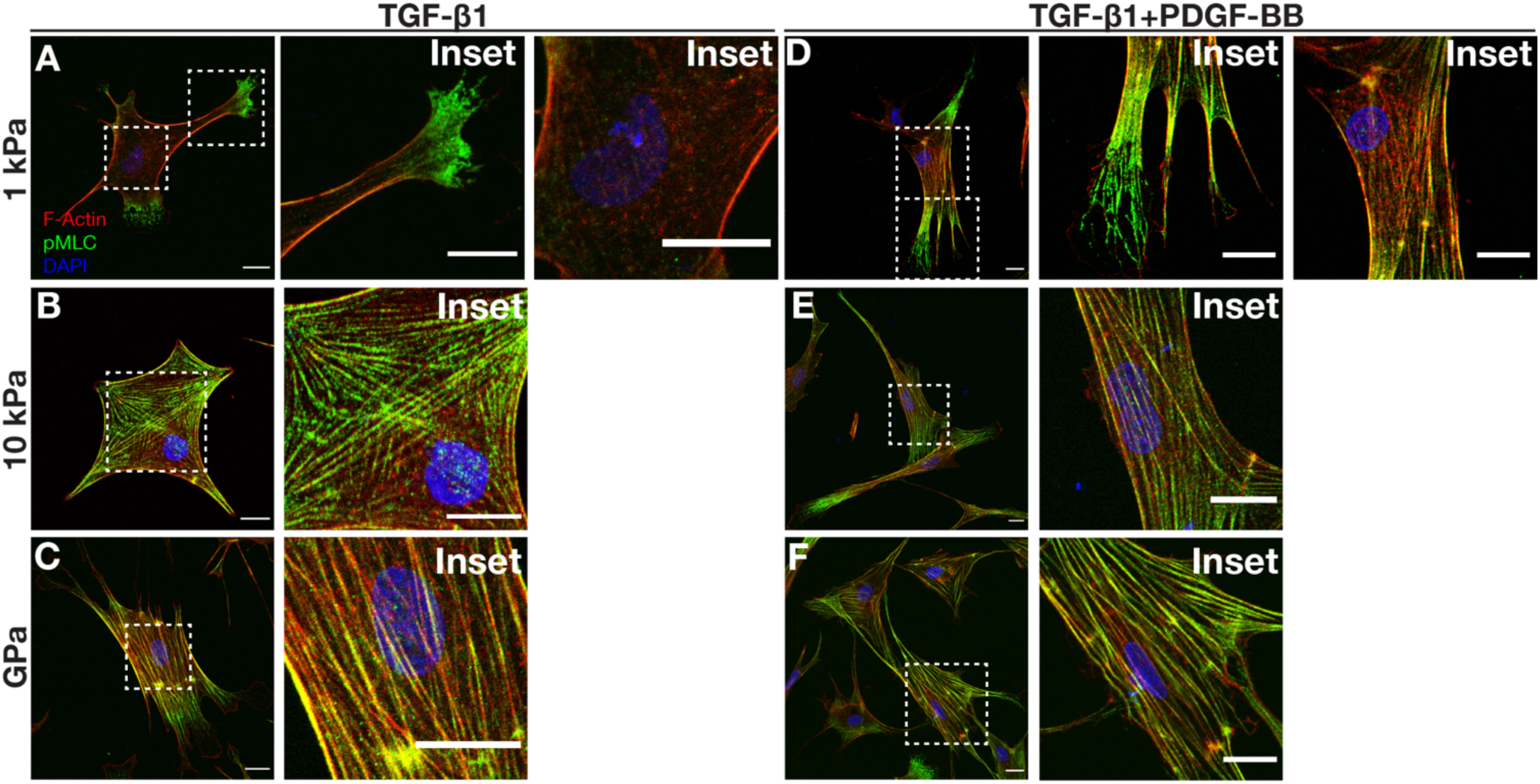
pMLC localizes to the tips of cell extensions on 1 kPa substrata in the presence of TGF-β1 and PDGF-BB. (A-F) Confocal images of phalloidin, DAPI, and pMLC immunofluorescence of corneal keratocytes in the presence of (A) (C) (E) TGF-β1 and (B) (D) (F) TGF-β1 and PDGF-BB. Focused insets of certain region of the cell are shown on the right panel. Scale bar = 20 µm. Error bars represent mean ± s.d. for n=6 substrates from 3 experimental replicates.

### Stiffness-dependent differences in FA patterning are unaffected by treatment with TGF-β1 and PDGF-BB

We have shown previously that stiffness-dependent differences in myofibroblast differentiation are regulated by signaling downstream of focal adhesions (28) and are associated with subcellular changes in FA patterning (28). Briefly, on stiff susbtrata, TGF-β1-treated keratocytes exhibit larger FAs, which are situated at the tips of stress fibers, while in a compliant microenvironment, the cells have smaller and fewer FAs, which are confined principally to the tips of cell extensions (28). To determine if treatment with both TGF-β1 and PDGF-BB alters FA patterning, we stained cells for vinculin immunofluorescence and quantified the FA size, number, and subcellular localization. In the serum-free and PDGF-BB treatment conditions, the cells had small FAs localized at the tips of cell extension, consistent with previous observations (28) (Fig. 5A-B). When treated with TGF-β1, as expected, we observed stiffness-dependent differences in FA patterning, as described above (Fig. 5C). Somewhat surprisingly, however, in the presence of both TGF-β1 and PDGF-BB, we observed the same subcellular distributions of FAs as those seen with TGF-β1 alone (Fig. 5C-D). On soft PA gels, the FAs were smaller and fewer in number than those observed within cells on either stiff PA substrata or collagen-coated glass coverslips (Fig. 5E-G, Fig. S2), even though the NRKs exhibited similar levels of ⍺-SMA staining in all these treatment conditions (Fig. 1D-E). Treatment with TGF-β1 and PDGF-BB does not therefore seem to alter stiffness-dependent differences in the subcellular patterning of FAs.

**Figure 5.**
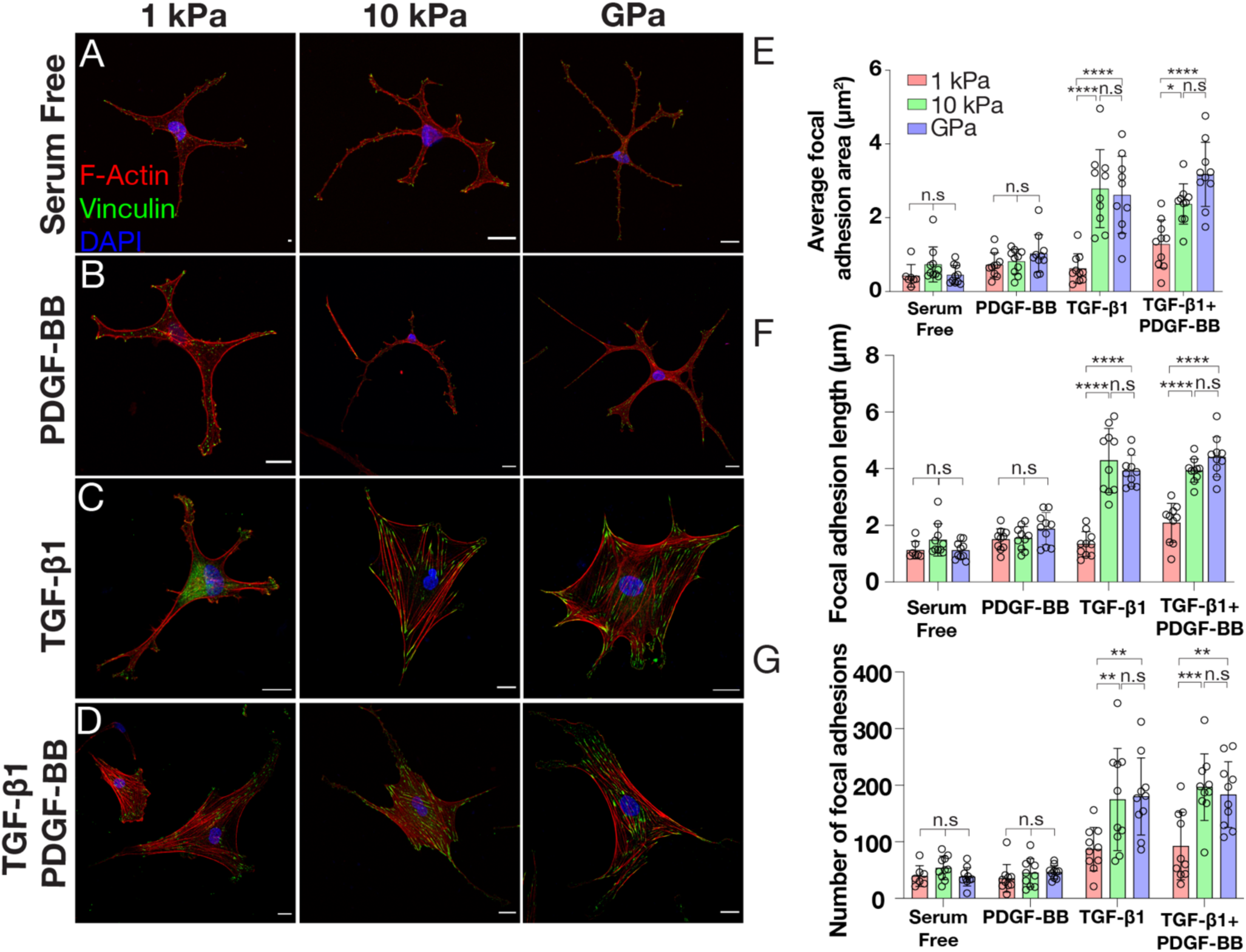
Subcellular localization of focal adhesions is regulated by substratum stiffness in the presence of TGF-β1 and PDGF-BB. (A–D) Confocal images of phalloidin, vinculin and DAPI immunofluorescence of corneal keratocytes in either (A) serum free conditions or in the presence of (B) PDGF-BB (C) TGF-β1 or (D) TGF-β1 and PDGF-BB. Scale bar = 20 µm. (E-G) Quantification of the (E) average focal adhesion (FA) area (F) FA length and (G) number of FA per cell. Error bars represent mean ± s.d. for n = 10 cells from 4 experimental replicates. A two-way ANOVA with a Tukey post-hoc test was used to evaluate significance among groups (*, p < 0.05; ****, p < 0.0001).

Taken together, our data indicate that the combined effects of TGF-β1 and PDGF-BB disrupt the stiffness-dependent myofibroblast differentiation of corneal keratocytes, decoupling the expression of myofibroblast markers from the elevated contractile phenotype typically associated with these cells. These results suggest potential crosstalk in the mechanotransductive signaling pathways downstream of TGF-β1 and PDGF-BB.

## Discussion

In a variety of fibroblastic cell types, changes in the mechanical properties of the ECM have been shown to influence myofibroblast differentiation (44–50). In the cornea, the tissue stiffens substantially following an injury (22, 23), and numerous cytokines, including TGF-β1 and PDGF-BB, are released into the corneal stroma (13–15). These change in tissue stiffness are associated with wound healing in the cornea and are thought to impact the behavior of corneal keratocytes, following an injury (22, 23, 51–54). Indeed, increased substratum stiffness has been shown to promote TGF-β1-mediated myofibroblast differentiation of cultured primary keratocytes where the cells exhibit elevated α-SMA immunofluorescence and exert higher traction stresses on stiffer substrata and this stiffness-dependent behavior depends on force-dependent signaling downstream of focal adhesions (28). In the presence of PDGF-BB, cultured NRKs exhibit stiffness-dependent differences in proliferation and morphology (27). Growth factor signaling and ECM stiffness impact keratocyte behavior, however it is unclear how signaling downstream of a combination of growth factors influence stiffness-dependent keratocyte response. PDGF signaling has also been shown to influence myofibroblast differentiation (20, 21, 40), but it is not known how treatment with PDGF affects stiffness-dependent differences in myofibroblast differentiation. Here, we cultured primary corneal keratocytes on polyacrylamide (PA) gels of different stiffnesses in the presence of both TGF-β1 and PDGF-BB to determine how combined biophysical and biochemical cues affect myofibroblast differentiation.

Interestingly, our results indicate that PDGF-BB disrupts stiffness-dependent differences in α-SMA staining in TGF-β1-treated keratocytes; however, it does not alter the effect of stiffness on cell contractility. In the presence of TGF-β1 alone, keratocytes on compliant environments exhibit decreased α-SMA immunofluorescence and contractility. When treated with both growth factors however, the cells formed prominent stress fibers and showed higher levels of α-SMA staining, even on soft substrata, although the cells did not exert elevated traction forces or mature focal adhesions (Fig. 6). Taken together, these data suggest that the presence of a combination of growth factors is enough to trigger the development of stress fibers and the expression of α-SMA. However, the creation of fully mature focal adhesions, achieved through integrin clustering, relies on the stiffness of the substrate. Without these adhesions, enhanced tractional forces cannot be generated.

**Figure 6.**
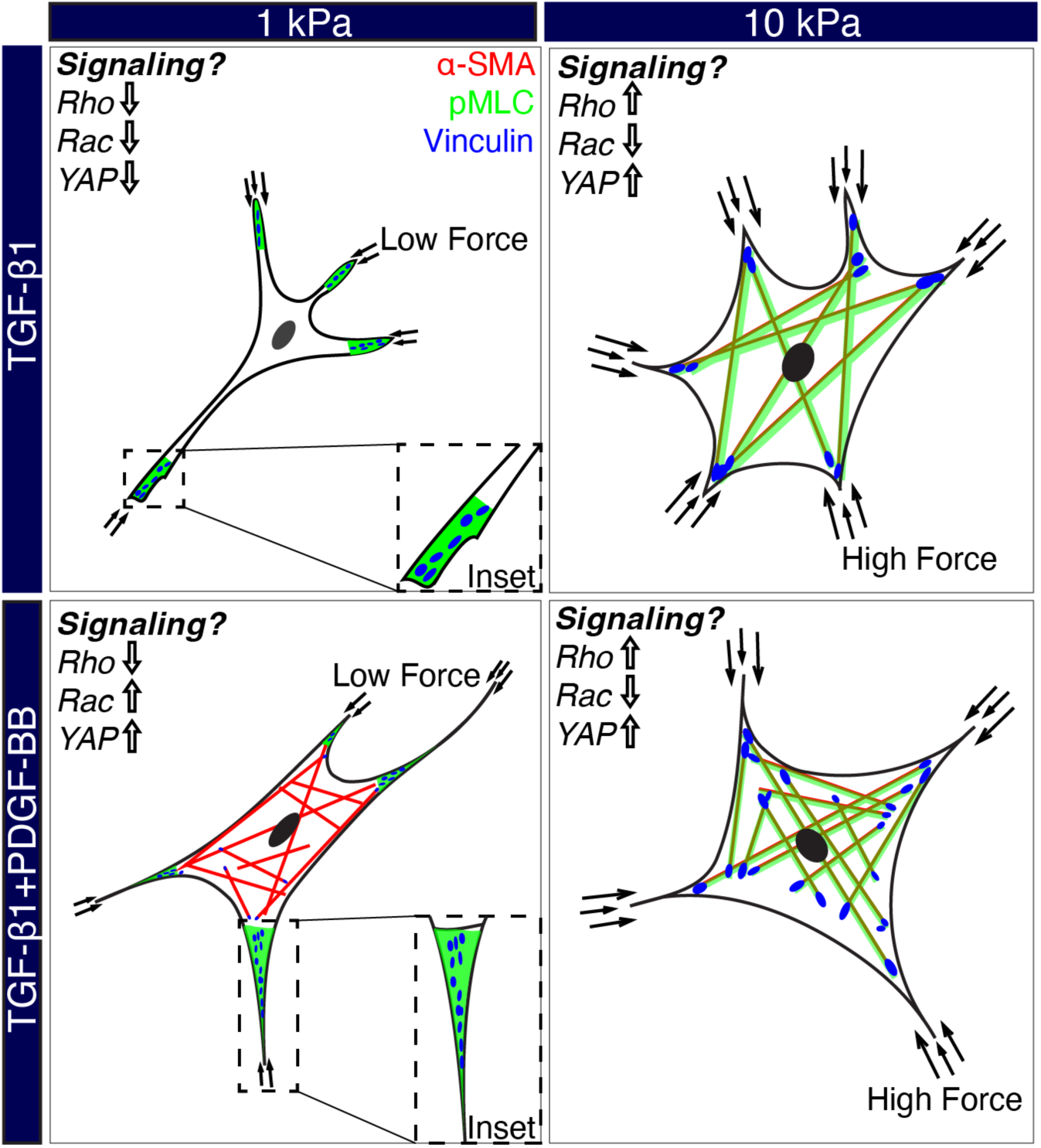
Schematic summarizing α-SMA expression, pMLC localization, subcellular patterning of focal adhesions and traction forces on keratocytes in the presence of TGF-β1 alone or in the presence of both TGF-β1 and PDGF-BB.

Somewhat surprisingly, our data suggest that treatment with PDGF-BB decouples force generation from TGF-β1 induced α-SMA expression. Myofibroblast differentiation is typically characterized by increased α-SMA expression, stress fiber formation, elevated cell contractile force and the formation of mature focal adhesions (25, 28, 42, 55–62). Previous studies in various cell types have linked increased traction force generation with elevated levels of α-SMA expression (41, 42, 61, 63, 64). Consistent with this idea, elevated tension promotes increased α-SMA expression and contractility in-vivo (43). Myofibroblasts also secrete a variety of ECM proteins that increases the stiffness of the surrounding matrix (12, 18, 65) . The cells are thought to counteract this change in mechanical properties by exerting larger traction stresses and forming more mature focal adhesions (63). Since treatment with TGF-β1 induces α-SMA expression and results in the formation of larger focal adhesions, a mechanical feedback loop which depends on α-SMA-containing stress fibers, is thought to regulate myofibroblast differentiation (56). In the presence of TGF-β1 alone, cultured corneal keratocytes exhibit stiffness-dependent differences in α-SMA expression, traction forces, and focal adhesion patterning that are in accord with this idea. However, when both TGF-β1 and PDGF-BB are present, we observed elevated α-SMA expression and stress fiber formation without a corresponding increase in cellular contractility or focal adhesion maturation. In soft substrata cultured with both TGF-β1 and PDGF-BB, contractility is mainly localized to the tips of the cell edges, as indicated by pMLC immunofluorescence, rather than along the full length of the cell’s actin stress fibers. This suggests that cells in this environment may not encounter enough mechanical resistance from the soft substrate to generate significant forces which lead to pMLC colocalization with stress fibers or the formation of mature focal adhesions. The development of mature focal adhesions is generally known to be a tension-dependent process (56, 63) and we hypothesize that sufficient tension cannot be developed on soft substrata to support the development of large focal adhesions. This may in turn reduce signaling downstream of focal adhesions which has been shown to modulate pMLC activity via Rho kinase and other mediators (66, 67). Our data thus suggests a nuanced relationship where α-SMA expression can be decoupled from myofibroblast differentiation. While these cells express α-SMA and form stress fibers, they do not manifest the full myofibroblast phenotype. Our discovery thus challenges conventional assumptions and underscores the complexity of cell behavior in response to growth factor signaling.

PDGF signaling has been shown to be important for myofibroblast differentiation in various fibrosis models, including the lungs (68) and kidneys (69) and blocking PDGF signaling inhibited corneal myofibroblast activation in other animal models (20, 40). Previous work in-vitro has shown that PDGF signaling is required for the TGF-β1-mediated myofibroblast differentiation of corneal keratocytes, specifically (21). Using function blocking antibodies to PDGF-AB ligand inhibited α-SMA expression in corneal keratocytes on glass substrata. Our experiments using pharmacological inhibition of PDGFR are consistent with these findings yet also suggest that PDGF signaling is important in mediating stiffness-dependent phenotypes in TGF-β1-induced myofibroblast differentiation. It has been suggested that the effect of PDGF is mediated via autocrine signaling (21, 70, 71), but it has not been shown which PDGF subunits are expressed in response to TGF-β1 nor how these might be altered by changes in substratum stiffness. Although previous studies have reported the concentrations of different growth factors present in human tear fluid during corneal wound healing (35, 72), the concentration of PDGF-BB secreted via autocrine signaling and how that compares to our experiments is still unknown. Our findings highlight the significance of signaling downstream of PDGF-BB in the TGF-β1-mediated myofibroblast differentiation. Although this suggests crosstalk between the pathways downstream of TGF-β1 and PDGF-BB, where these pathways interact is not fully understood. There are numerous signaling molecules within both cascades, such as PI3-kinase and Akt among others (36, 73–75), which may be points of crosstalk. The Rho family of small GTPases for instance, comprising Rho, Rac, and Cdc42, play a crucial role in regulating cell mechanical activity in response to growth factors and cytokines across various cell types (49, 76–78). In fibroblasts, activated Rho promotes stress fiber formation and focal adhesion development (79–81), and Rho-mediated signaling is known to play a key role in TGF-β1 -induced α-SMA expression and contractility (26, 82, 83). Conversely, Rac activation by PDGF BB induces cell elongation and rapid migration in corneal and dermal fibroblasts (84, 85). Since keratocyte growth factor responses appear to be regulated by the interplay between Rho and Rac signaling, and the structural and mechanical properties of the ECM (86), in our experiments with both TGF-β1 and PDGF-BB, we speculate that on stiffer substrata, Rho-mediated signaling is activated, evident by highly contractile cells forming stress fibers and exerting elevated traction stresses. In softer conditions however, where keratocytes exhibited elevated α-SMA staining without a proportional increase in contractility, we speculate increased Rac activation but not Rho (Fig. 6). Additionally, exploring the downstream events in the TGF-β1 signaling cascade following TGF-β1 receptor activation may shed light on whether SMAD activity is influenced by the presence of both growth factors on substrata of varying stiffnesses (28, 87–89). It would also be interesting to study the transcriptional changes associated with corneal keratocyte behavior when exposed to different soluble cues. This approach offers an unbiased exploration of the gene expression profile, potentially elucidating the molecular basis for the observed phenotypic changes. Identifying relevant targets within the signaling cascade through these transcriptional studies could guide future experiments, paving the way for a more comprehensive understanding of myofibroblast differentiation and its modulation in tissue repair and fibrosis management.

In conclusion, our study not only uncovers surprising nuances in keratocyte behavior but also highlights the need for further investigation into the intricate signaling pathways governing myofibroblast differentiation. The ability to decouple α-SMA expression from other myofibroblast associated behavior opens up exciting prospects for more targeted investigations in tissue repair and fibrotic conditions. As we continue to unravel the complexities of these processes, the potential for innovative therapeutic strategies becomes increasingly promising.

## Author contributions

KSI, VDV, WMP, and DWS conceived the study and designed experiments. KSI and DPM conducted all experiments and analyzed all experimental data. All authors discussed and interpreted results. KSI and VDV wrote the manuscript with feedback from all authors.

## Funding

This work was supported by NIH Grants R01 EY030190 and P30 EY030413, and a Challenge Grant from Research to Prevent Blindness.

## Acknowledgments

We would also like to thank the members of the Varner, Petroll, and Schmidtke labs for their many helpful discussions and comments.

---

**Figure S1.**
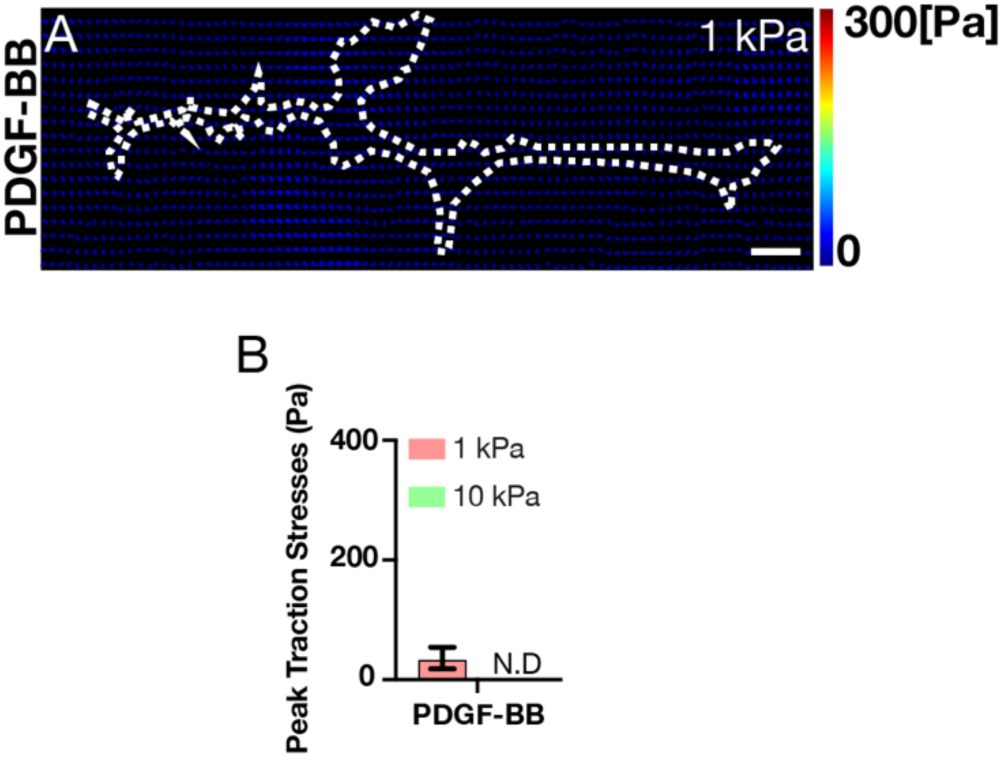
The traction stresses exerted by PDGF-BB treated keratocytes is minimal on soft substrata and in the not detectable (N.D) range on stiff substrata. (A) Computed traction stresses for representative NRKs cultured on 1 kPa substrata in the presence of PDGF-BB. (B) Quantification of peak traction stress after 5 days of culture in the presence of growth factor. Error bars represent mean ± s.d. for individual cells from 4 experimental replicates. A two-way ANOVA with a Tukey post-hoc test was used to evaluate significance among groups (*, p < 0.05; ****, p < 0.0001).

**Figure S2.**
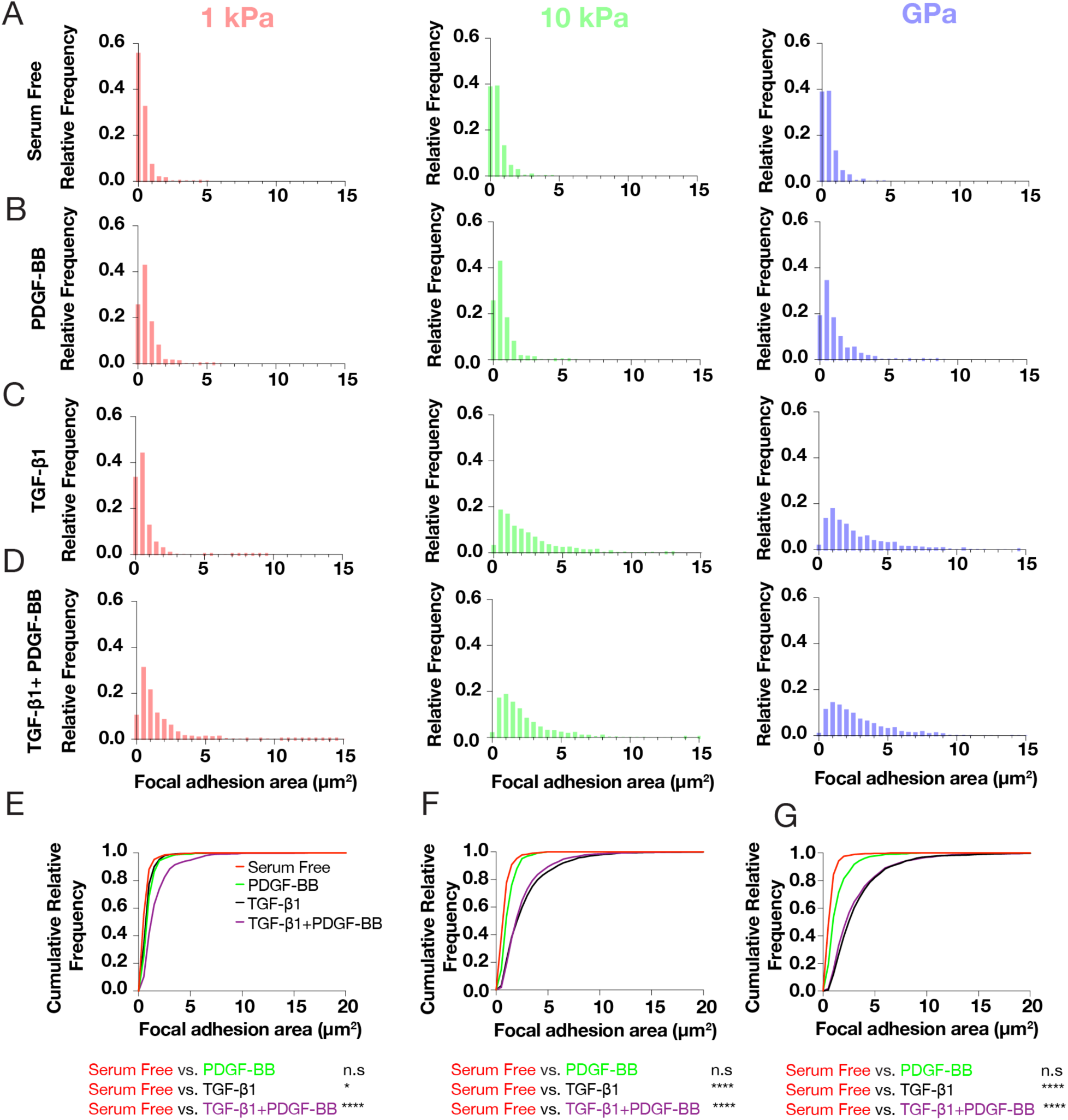
Focal adhesion area is regulated by substratum stiffness in the presence of TGF-β1 and PDGF-BB. (A-D) Relative frequency histograms of FA area in either (A) serum free conditions or in the presence of (B) PDGF-BB (C) TGF-β1 or (D) TGF-β1 and PDGF-BB. (E-G). Cumulative frequency plots were compared statistically using a Kolmogorov-Smirnov test.

## Notes

### Competing Interest Statement

The authors have declared no competing interest.

## References

1. Fini, M.E. 1999. Keratocyte and fibroblast phenotypes in the repairing cornea. Prog Retin Eye Res. 18:529–551.

2. Meek, K.M., and C. Boote. 2009. The use of X-ray scattering techniques to quantify the orientation and distribution of collagen in the corneal stroma. Prog Retin Eye Res. 28:369–392.

3. Meek, K.M. 2009. Corneal collagen—its role in maintaining corneal shape and transparency. Biophysical Rev. 1:83–93.

4. Nishida, T., K. Yasumoto, T. Otori, and J. Desaki. 1988. The network structure of corneal fibroblasts in the rat as revealed by scanning electron microscopy. Invest Ophth Vis Sci. 29:1887–90.

5. Hogan, M.J., J.A. Alvarado, and J. E. Weddell. 1971. Histology of the Human Eye. An atlas and textbook.

6. Hassell, J.R., and D.E. Birk. 2010. The molecular basis of corneal transparency. Exp Eye Res. 91:326–335.

7. Beales, M.P., J.L. Funderburgh, J.V. Jester, and J.R. Hassell. 1999. Proteoglycan synthesis by bovine keratocytes and corneal fibroblasts: maintenance of the keratocyte phenotype in culture. Invest Ophth Vis Sci. 40:1658–63.

8. Jester, J.V., P.A. Barry, G.J. Lind, W.M. Petroll, R. Garana, and H.D. Cavanagh. 1994. Corneal keratocytes: in situ and in vitro organization of cytoskeletal contractile proteins. Invest Ophth Vis Sci. 35:730–43.

9. Zieske, J.D., S.R. Guimarães, and A.E.K. Hutcheon. 2001. Kinetics of Keratocyte Proliferation in Response to Epithelial Debridement. Exp Eye Res. 72:33–39.

10. Boote, C., Y. Du, S. Morgan, J. Harris, C.S. Kamma-Lorger, S. Hayes, K.L. Lathrop, D.S. Roh, M.K. Burrow, J. Hiller, N.J. Terrill, J.L. Funderburgh, and K.M. Meek. 2012. Quantitative Assessment of Ultrastructure and Light Scatter in Mouse Corneal Debridement WoundsUltrastructure and Light Scatter in Corneal Wounds. Invest Ophth Vis Sci. 53:2786– 2795.

11. Singh, V., V. Agrawal, M.R. Santhiago, and S.E. Wilson. 2012. Stromal fibroblast–bone marrow-derived cell interactions: Implications for myofibroblast development in the cornea. Exp Eye Res. 98:1–8.

12. Chen, C., B. Michelini-Norris, S. Stevens, J. Rowsey, X. Ren, M. Goldstein, and G. Schultz. 2000. Measurement of mRNAs for TGFss and extracellular matrix proteins in corneas of rats after PRK. Investig. Ophthalmol. Vis. Sci. 41:4108–16.

13. Imanishi, J., K. Kamiyama, I. Iguchi, M. Kita, C. Sotozono, and S. Kinoshita. 2000. Growth factors: importance in wound healing and maintenance of transparency of the cornea. Prog Retin Eye Res. 19:113–129.

14. Torricelli, A.A.M., V. Singh, M.R. Santhiago, and S.E. Wilson. 2013. The Corneal Epithelial Basement Membrane: Structure, Function, and DiseaseCorneal Basement Membrane. Invest Ophth Vis Sci. 54:6390–6400.

15. Wilson, S.E., R.R. Mohan, R.R. Mohan, R. Ambrósio, J. Hong, and J. Lee. 2001. The Corneal Wound Healing Response: Cytokine-mediated Interaction of the Epithelium, Stroma, and Inflammatory Cells. Prog Retin Eye Res. 20:625–637.

16. Kurosaka, H., D. Kurosaka, K. Kato, Y. Mashima, and Y. Tanaka. 1998. Transforming growth factor-beta 1 promotes contraction of collagen gel by bovine corneal fibroblasts through differentiation of myofibroblasts. Invest Ophth Vis Sci. 39:699–704.

17. Nakamura, K. 2003. Interaction Between Injured Corneal Epithelial Cells and Stromal Cells. Cornea. 22:S35–S47.

18. Jester, J.V., J. Huang, P.A. Barry-Lane, W.W. Kao, W.M. Petroll, and H.D. Cavanagh. 1999. Transforming growth factor(beta)-mediated corneal myofibroblast differentiation requires actin and fibronectin assembly. Invest Ophth Vis Sci. 40:1959–67.

19. Kim, A., N. Lakshman, D. Karamichos, and W.M. Petroll. 2009. Growth factor regulation of corneal keratocyte differentiation and migration in compressed collagen matrices. Invest Ophth Vis Sci. 51:864–75.

20. Kaur, H., S.S. Chaurasia, F.W. de Medeiros, V. Agrawal, M.Q. Salomao, N. Singh, B.K. Ambati, and S.E. Wilson. 2009. Corneal stroma PDGF blockade and myofibroblast development. Exp. Eye Res. 88:960–965.

21. Jester, J.V., J. Huang, W.M. Petroll, and H.D. Cavanagh. 2002. TGFβ Induced Myofibroblast Differentiation of Rabbit Keratocytes Requires Synergistic TGFβ, PDGF and Integrin Signaling. Exp Eye Res. 75:645–657.

22. Thomasy, S.M., V.K. Raghunathan, M. Winkler, C.M. Reilly, A.R. Sadeli, P. Russell, J.V. Jester, and C.J. Murphy. 2014. Elastic modulus and collagen organization of the rabbit cornea: Epithelium to endothelium. Acta Biomater. 10:785–791.

23. Raghunathan, V.K., S.M. Thomasy, P. Strøm, B. Yañez-Soto, S.P. Garland, J. Sermeno, C.M. Reilly, and C.J. Murphy. 2017. Tissue and cellular biomechanics during corneal wound injury and repair. Acta Biomater. 58:291–301.

24. Dreier, B., S.M. Thomasy, R. Mendonsa, V.K. Raghunathan, P. Russell, and C.J. Murphy. 2013. Substratum Compliance Modulates Corneal Fibroblast to Myofibroblast Transformation. Investigative Opthalmology Vis Sci. 54:5901.

25. Maruri, D.P., M. Miron-Mendoza, P.B. Kivanany, J.M. Hack, D.W. Schmidtke, W.M. Petroll, and V.D. Varner. 2020. ECM Stiffness Controls the Activation and Contractility of Corneal Keratocytes in Response to TGF-β1. Biophys J. 119:1865–1877.

26. Lakshman, N., and W.M. Petroll. 2012. Growth factor regulation of corneal keratocyte mechanical phenotypes in 3-D collagen matrices. Invest Ophth Vis Sci. 53:1077–86.

27. Iyer, K.S., D.P. Maruri, K.E. Peak, D.W. Schmidtke, W.M. Petroll, and V.D. Varner. 2022. ECM stiffness modulates the proliferation but not the motility of primary corneal keratocytes in response to PDGF-BB. Exp Eye Res. 220:109112.

28. Maruri, D.P., K.S. Iyer, D.W. Schmidtke, W.M. Petroll, and V.D. Varner. 2022. Signaling Downstream of Focal Adhesions Regulates Stiffness-Dependent Differences in the TGF-β1-Mediated Myofibroblast Differentiation of Corneal Keratocytes. Frontiers Cell Dev Biology. 10:886759.

29. Kim, S., I. Jalilian, S.M. Thomasy, M.A.W. Bowman, V.K. Raghunathan, Y. Song, C.A. Reinhart-King, and C.J. Murphy. 2020. Intrastromal Injection of Hyaluronidase Alters the Structural and Biomechanical Properties of the Corneal Stroma. Transl Vis Sci Technology. 9:21.

30. Miron-Mendoza, M., E. Graham, P. Kivanany, J. Quiring, and W.M. Petroll. 2015. The Role of Thrombin and Cell Contractility in Regulating Clustering and Collective Migration of Corneal Fibroblasts in Different ECM Environments. Invest Ophth Vis Sci. 56:2079–90.

31. Kivanany, P.B., K.C. Grose, N. Yonet-Tanyeri, S. Manohar, Y. Sunkara, K.H. Lam, D.W. Schmidtke, V.D. Varner, and W.M. Petroll. 2018. An In Vitro Model for Assessing Corneal Keratocyte Spreading and Migration on Aligned Fibrillar Collagen. J Funct Biomaterials. 9:54.

32. Jester, J.V., P.A. Barry-Lane, H.D. Cavanagh, and W.M. Petroll. 1996. Induction of α-Smooth Muscle Actin Expression and Myofibroblast Transformation in Cultured Corneal Keratocytes. Cornea. 15:505–516.

33. Jester, J.V., and J. Ho-Chang. 2003. Modulation of cultured corneal keratocyte phenotype by growth factors/cytokines control in vitro contractility and extracellular matrix contraction. Exp Eye Res. 77:581–592.

34. Lakshman, N., A. Kim, and W.M. Petroll. 2010. Characterization of corneal keratocyte morphology and mechanical activity within 3-D collagen matrices. Exp Eye Res. 90:350–359.

35. Hoppenreijs, V.P., E. Pels, G.F. Vrensen, P.C. Felten, and W.F. Treffers. 1993. Platelet-derived growth factor: receptor expression in corneas and effects on corneal cells. Investig. Ophthalmol. Vis. Sci. 34:637–49.

36. Dell, S., S. Peters, P. Müther, N. Kociok, and A.M. Joussen. 2006. The role of PDGF receptor inhibitors and PI3-kinase signaling in the pathogenesis of corneal neovascularization. Invest Ophth Vis Sci. 47:1928–37.

37. Martiel, J.-L., A. Leal, L. Kurzawa, M. Balland, I. Wang, T. Vignaud, Q. Tseng, and M. Théry. 2015. Measurement of cell traction forces with ImageJ. Methods Cell Biol. 125:269–87.

38. Maruthamuthu, V., B. Sabass, U.S. Schwarz, and M.L. Gardel. 2011. Cell-ECM traction force modulates endogenous tension at cell–cell contacts. Proc National Acad Sci. 108:4708– 4713.

39. Kivanany, P.B., K.C. Grose, N. Yonet-Tanyeri, S. Manohar, Y. Sunkara, K.H. Lam, D.W. Schmidtke, V.D. Varner, and W.M. Petroll. 2018. An In Vitro Model for Assessing Corneal Keratocyte Spreading and Migration on Aligned Fibrillar Collagen. J Funct Biomaterials. 9:54.

40. Singh, V., M.R. Santhiago, F.L. Barbosa, V. Agrawal, N. Singh, B.K. Ambati, and S.E. Wilson. 2011. Effect of TGFβ and PDGF-B blockade on corneal myofibroblast development in mice. Exp. Eye Res. 93:810–817.

41. Chen, J., H. Li, N. SundarRaj, and J.H. -C. Wang. 2007. Alpha-smooth muscle actin expression enhances cell traction force. Cell Motil. Cytoskeleton. 64:248–257.

42. Hinz, B., G. Celetta, J.J. Tomasek, G. Gabbiani, and C. Chaponnier. 2001. Alpha-Smooth Muscle Actin Expression Upregulates Fibroblast Contractile Activity. Mol. Biol. Cell. 12:2730– 2741.

43. Hinz, B., D. Mastrangelo, C.E. Iselin, C. Chaponnier, and G. Gabbiani. 2001. Mechanical Tension Controls Granulation Tissue Contractile Activity and Myofibroblast Differentiation. Am. J. Pathol. 159:1009–1020.

44. Hinz, B., S.H. Phan, V.J. Thannickal, A. Galli, M.-L. Bochaton-Piallat, and G. Gabbiani. 2007. The Myofibroblast. Am J Pathology. 170:1807–1816.

45. Grinnell, F., and W.M. Petroll. 2010. Cell Motility and Mechanics in Three-Dimensional Collagen Matrices. Annu Rev Cell Dev Bi. 26:335–361.

46. Klingberg, F., B. Hinz, and E.S. White. 2013. The myofibroblast matrix: implications for tissue repair and fibrosis. J. Pathol. 229:298–309.

47. Wells, R.G. 2013. Tissue mechanics and fibrosis. Biochimica Et Biophysica Acta Bba - Mol Basis Dis. 1832:884–890.

48. Falke, L.L., S. Gholizadeh, R. Goldschmeding, R.J. Kok, and T.Q. Nguyen. 2015. Diverse origins of the myofibroblast—implications for kidney fibrosis. Nat Rev Nephrol. 11:233–244.

49. Petroll, W.M., and M. Miron-Mendoza. 2015. Mechanical interactions and crosstalk between corneal keratocytes and the extracellular matrix. Exp Eye Res. 133:49–57.

50. Zhou, Y., J.C. Horowitz, A. Naba, N. Ambalavanan, K. Atabai, J. Balestrini, P.B. Bitterman, R.A. Corley, B.-S. Ding, A.J. Engler, K.C. Hansen, J.S. Hagood, F. Kheradmand, Q.S. Lin, E. Neptune, L. Niklason, L.A. Ortiz, W.C. Parks, D.J. Tschumperlin, E.S. White, H.A. Chapman, and V.J. Thannickal. 2018. Extracellular matrix in lung development, homeostasis and disease. Matrix Biol. 73:77–104.

51. Kim, A., C. Zhou, N. Lakshman, and W.M. Petroll. 2012. Corneal stromal cells use both high- and low-contractility migration mechanisms in 3-D collagen matrices. Exp. Cell Res. 318:741–752.

52. Grinnell, F., and W.M. Petroll. 2010. Cell Motility and Mechanics in Three-Dimensional Collagen Matrices. Annu. Rev. cell Dev. Biol. 26:335–361.

53. Karamichos, D., N. Lakshman, and W.M. Petroll. 2009. An experimental model for assessing fibroblast migration in 3-D collagen matrices. Cell Motil. Cytoskelet. 66:1–9.

54. Petroll, W.M., V.D. Varner, and D.W. Schmidtke. 2020. Keratocyte mechanobiology. Exp Eye Res. 200:108228.

55. Hinz, B. 2016. Myofibroblasts. Exp Eye Res. 142:56–70.

56. Hinz, B. 2007. Formation and Function of the Myofibroblast during Tissue Repair. J Invest Dermatol. 127:526–537.

57. Hinz, B. 2010. The myofibroblast: Paradigm for a mechanically active cell. J Biomech. 43:146–155.

58. Jester, J.V., and J. Ho-Chang. 2003. Modulation of cultured corneal keratocyte phenotype by growth factors/cytokines control in vitro contractility and extracellular matrix contraction. Exp. Eye Res. 77:581–592.

59. Maruri, D.P., K.S. Iyer, D.W. Schmidtke, W.M. Petroll, and V.D. Varner. 2022. Signaling Downstream of Focal Adhesions Regulates Stiffness-Dependent Differences in the TGF-β1-Mediated Myofibroblast Differentiation of Corneal Keratocytes. Front. Cell Dev. Biol. 10:886759.

60. Petroll, W.M., L. Ma, and J.V. Jester. 2003. Direct correlation of collagen matrix deformation with focal adhesion dynamics in living corneal fibroblasts. J. Cell Sci. 116:1481–1491.

61. Hinz, B., and G. Gabbiani. 2003. Mechanisms of force generation and transmission by myofibroblasts. Curr. Opin. Biotechnol. 14:538–546.

62. Malmström, J., H. Lindberg, C. Lindberg, C. Bratt, E. Wieslander, E.-L. Delander, B. Särnstrand, J.S. Burns, P. Mose-Larsen, S. Fey, and G. Marko-Varga. 2004. Transforming Growth Factor-β1 Specifically Induce Proteins Involved in the Myofibroblast Contractile Apparatus*. Mol. Cell. Proteom. 3:466–477.

63. Hinz, B., V. Dugina, C. Ballestrem, B. Wehrle-Haller, and C. Chaponnier. 2003. α-Smooth Muscle Actin Is Crucial for Focal Adhesion Maturation in Myofibroblasts. Mol. Biol. Cell. 14:2508–2519.

64. Wrobel, L.K., T.R. Fray, J.E. Molloy, J.J. Adams, M.P. Armitage, and J.C. Sparrow. 2002. Contractility of single human dermal myofibroblasts and fibroblasts. Cell Motil. Cytoskelet. 52:82–90.

65. Blalock, T.D., M.R. Duncan, J.C. Varela, M.H. Goldstein, S.S. Tuli, G.R. Grotendorst, and G.S. Schultz. 2003. Connective Tissue Growth Factor Expression and Action in Human Corneal Fibroblast Cultures and Rat Corneas after Photorefractive Keratectomy. Investig. Opthalmology Vis. Sci. 44:1879.

66. Schober, M., S. Raghavan, M. Nikolova, L. Polak, H.A. Pasolli, H.E. Beggs, L.F. Reichardt, and E. Fuchs. 2007. Focal adhesion kinase modulates tension signaling to control actin and focal adhesion dynamics. J. Cell Biol. 176:667–680.

67. Fan, L., A. Sebe, Z. Péterfi, A. Masszi, A.C.P. Thirone, O.D. Rotstein, H. Nakano, C.A. McCulloch, K. Szászi, I. Mucsi, and A. Kapus. 2007. Cell Contact–dependent Regulation of Epithelial–Myofibroblast Transition via the Rho-Rho Kinase-Phospho-Myosin Pathway. Mol. Biol. Cell. 18:1083–1097.

68. Wollin, L., I. Maillet, V. Quesniaux, A. Holweg, and B. Ryffel. 2014. Antifibrotic and Anti-inflammatory Activity of the Tyrosine Kinase Inhibitor Nintedanib in Experimental Models of Lung Fibrosis. J. Pharmacol. Exp. Ther. 349:209–220.

69. Chen, Y.-T., F.-C. Chang, C.-F. Wu, Y.-H. Chou, H.-L. Hsu, W.-C. Chiang, J. Shen, Y.-M. Chen, K.-D. Wu, T.-J. Tsai, J.S. Duffield, and S.-L. Lin. 2011. Platelet-derived growth factor receptor signaling activates pericyte–myofibroblast transition in obstructive and post-ischemic kidney fibrosis. Kidney Int. 80:1170–1181.

70. Jester, J.V., J. Huang, S. Fisher, J. Spiekerman, J.H. Chang, W.E. Wright, and J.W. Shay. 2003. Myofibroblast Differentiation of Normal Human Keratocytes and hTERT, Extended-Life Human Corneal Fibroblasts. Investig. Opthalmology Vis. Sci. 44:1850.

71. Soma, Y., and G.R. Grotendorst. 1989. TGF-beta stimulates primary human skin fibroblast DNA synthesis via an autocrine production of PDGF-related peptides. J. Cell. Physiol. 140:246– 53.

72. Vesaluoma, M., A.-M. Teppo, C. Grönhagen-Riska, and T. Tervo. 1997. Platelet-derived growth factor-BB (PDGF-BB) in tear fluid: a potential modulator of corneal wound healing following photorefractive keratectomy. Curr. Eye Res. 16:825–831.

73. Conte, E., M. Fruciano, E. Fagone, E. Gili, F. Caraci, M. Iemmolo, N. Crimi, and C. Vancheri. 2011. Inhibition of PI3K Prevents the Proliferation and Differentiation of Human Lung Fibroblasts into Myofibroblasts: The Role of Class I P110 Isoforms. PLoS ONE. 6:e24663.

74. Wilkes, M.C., H. Mitchell, S.G. Penheiter, J.J. Doré, K. Suzuki, M. Edens, D.K. Sharma, R.E. Pagano, and E.B. Leof. 2005. Transforming Growth Factor-β Activation of Phosphatidylinositol 3-Kinase Is Independent of Smad2 and Smad3 and Regulates Fibroblast Responses via p21-Activated Kinase-2. Cancer Res. 65:10431–10440.

75. Hettiarachchi, S.U., Y.-H. Li, J. Roy, F. Zhang, E. Puchulu-Campanella, S.D. Lindeman, M. Srinivasarao, K. Tsoyi, X. Liang, E.A. Ayaub, C. Nickerson-Nutter, I.O. Rosas, and P.S. Low. 2020. Targeted inhibition of PI3 kinase/mTOR specifically in fibrotic lung fibroblasts suppresses pulmonary fibrosis in experimental models. Sci. Transl. Med. 12.

76. Jaffe, A.B., and A. Hall. 2005. RHO GTPASES: Biochemistry and Biology. Cell Dev. Biol. 21:247–269.

77. Hall, A. 2005. Rho GTPases and the control of cell behaviour. Biochem. Soc. Trans. 33:891–895.

78. Wang, J., X. Liu, and Y. Zhong. 2013. Rho/Rho-associated kinase pathway in glaucoma (Review). Int. J. Oncol. 43:1357–67.

79. Anderson, S., L. DiCesare, I. Tan, T. Leung, and N. SundarRaj. 2004. Rho-mediated assembly of stress fibers is differentially regulated in corneal fibroblasts and myofibroblasts. Exp. Cell Res. 298:574–583.

80. Parizi, M., E.W. Howard, and J.J. Tomasek. 2000. Regulation of LPA-Promoted Myofibroblast Contraction: Role of Rho, Myosin Light Chain Kinase, and Myosin Light Chain Phosphatase. Exp. Cell Res. 254:210–220.

81. Totsukawa, G., Y. Yamakita, S. Yamashiro, D.J. Hartshorne, Y. Sasaki, and F. Matsumura. 2000. Distinct Roles of Rock (Rho-Kinase) and Mlck in Spatial Regulation of Mlc Phosphorylation for Assembly of Stress Fibers and Focal Adhesions in 3t3 Fibroblasts. J. Cell Biol. 150:797–806.

82. Chen, J., E. Guerriero, Y. Sado, and N. SundarRaj. 2009. Rho-mediated regulation of TGF-beta1- and FGF-2-induced activation of corneal stromal keratocytes. Investig. Ophthalmol. Vis. Sci. 50:3662–70.

83. Yamamoto, M., A.J. Quantock, R.D. Young, N. Okumura, M. Ueno, Y. Sakamoto, S. Kinoshita, and N. Koizumi. 2012. A selective inhibitor of the Rho kinase pathway, Y-27632, and its influence on wound healing in the corneal stroma. Mol. Vis. 18:1727–39.

84. Grinnell, F. 2000. Fibroblast–collagen-matrix contraction: growth-factor signalling and mechanical loading. Trends Cell Biol. 10:362–365.

85. Sander, E.E., J.P. ten Klooster, S. van Delft, R.A. van der Kammen, and J.G. Collard. 1999. Rac downregulates Rho activity: reciprocal balance between both GTPases determines cellular morphology and migratory behavior. J. cell Biol. 147:1009–22.

86. Petroll, W.M., and N. Lakshman. 2015. Fibroblastic Transformation of Corneal Keratocytes by Rac Inhibition is Modulated by Extracellular Matrix Structure and Stiffness. J Funct Biomaterials. 6:222–240.

87. Tang, Y., E. Du, G. Wang, F. Qin, Z. Meng, L. Dai, Y. Wang, and S. Ren. 2023. A negative feedback loop centered on SMAD3 expression in transforming growth factor β1-induced corneal myofibroblast differentiation. Exp. Eye Res. 236:109654.

88. Saika, S., O. Yamanaka, Y. Okada, and T. Sumioka. 2016. Modulation of Smad signaling by non-TGFβ components in myofibroblast generation during wound healing in corneal stroma. Exp. Eye Res. 142:40–48.

89. Saika, S. 2004. TGF-β Signal Transduction in Corneal Wound Healing as a Therapeutic Target. Cornea. 23:S25–S30.

